# VirProtRAG: Literature-grounded viral protein function annotation with retrieval-augmented generation

**DOI:** 10.64898/2026.07.03.736267

**Authors:** Jiaojiao Guan, Jiayu Shang, Cheng Peng, Yanni Sun

## Abstract

Viruses play indispensable roles in ecosystems and human health, yet deciphering their molecular functions remains challenging. Many viral protein annotations are incomplete or poorly characterized. Existing tools typically predict functional categories without linking to verifiable evidence, hindering the credibility of functional interpretation. Here, we present VirProtRAG, a viral protein function annotation framework that integrates information retrieval with evidence-grounded knowledge generation. It introduces three task-adapted components: a hybrid retrieval module combining keyword-based and semantic dense retrieval to maximize literature coverage, synonym-expanded and rank-aware retrieval with reciprocal rank fusion for improved search effectiveness, and literature quality and evidence-oriented re-ranking to enhance reliability and interpretability. Results show that hybrid retrieval strategy performed best, with quality and evidence features further enhancing re-ranking. Compared with direct LLM prompting without retrieved literature, it consistently improves generation performance, underscoring the critical role of external knowledge. Finally, we built a searchable database comprising all 17,484 reviewed Swiss-Prot viral proteins, supporting both sequence- and text-based queries. VirProtRAG introduced 32.53% non-overlapping function annotations beyond existing expert curation, and independently supported 56.34% of sequence-inferred function points with retrieved literature. Case studies further demonstrate its capability to augment and refine the characterization of previously unannotated or poorly understood viral proteins.

**Key Messages:**

- VirProtRAG bridges large-scale biomedical literature and viral protein annotation, enabling traceable and biologically interpretable annotations grounded in verifiable publications.
- By integrating hybrid sparse–dense retrieval, synonym expansion, and adaptive re-ranking using literature quality and evidence-type features, VirProtRAG substantially improves recall and ranking of biologically relevant publications, helping ground LLM outputs in traceable literature evidence.
- Leveraging this framework, VirProtRAG automatically annotated all 17,484 reviewed Swiss-Prot viral proteins under the Virus taxonomic group and established an interactive, continuously expandable database. The resource may assist curators in rapid validation while supporting large-scale studies of viral evolution and host interactions.

## Introduction

Viruses are the most abundant biological entities on Earth, with a ubiquitous presence across all known ecosystems. They not only include the major pathogens of human diseases [1, 2], from acute pandemics to persistent infections, but also play indispensable ecological roles, mediating horizontal gene transfer, shaping host evolution, and regulating microbial community dynamics throughout the biosphere [3, 4]. The study of viruses has long been central to understanding fundamental cellular processes and continues to inspire novel therapeutic and biotechnological innovations [5].

Despite the rapid accumulation of viral genomic data enabled by high-throughput sequencing technologies [6], their function annotation remains markedly lagging [7]. First, in the UniProtKB/Swiss-Prot database [8], a substantial fraction of viral proteins remains unannotated. In particular, 27.49% of viral entries contain only protein names but no functional descriptions. Second, even when annotations exist, they are often overly generic and provide limited mechanistic insight. Labels such as “nonstructural protein,” “VP2,” or “ORF1” primarily indicate genomic position or structural category and do not specify enzymatic activity, interaction partners, or mechanistic roles in the viral life cycle. Third, annotations are frequently incomplete [9]: assigned Gene Ontology (GO) terms [10] are often limited to high-level categories (e.g., GO:0005488, *binding*), whereas more specific descendant terms that capture precise biochemical functions remain missing. The reliance on manual expert curation is slow, labor-intensive, and difficult to scale with the exponential growth of sequence data.

The limited amount of curated and fine-grained functional data hinders robust model training and compromises generalization to newly sequenced or poorly characterized viral taxa. In addition, this scarcity of reliable annotations restricts our ability to formulate mechanistic hypotheses, interpret predictions in a biological context, and design targeted experiments to validate viral protein functions. In this regard, enriching and refining existing annotations is crucial for advancing both data-driven modeling and experimental discovery.

As research on viral proteins expands, a vast amount of mechanistic information has already been reported in the literature. Extracting and integrating this dispersed knowledge from publications represents a promising approach for systematic annotation enrichment and biological discovery [11, 12]. Recent advances in text-based annotation highlight this potential. For instance, GORetriever [13] formulates protein function prediction as a two-stage “retrieve-rerank” process: it first retrieves candidate GO terms and representative functional sentences from curated literature, then employs a cross-encoder to perform deep semantic interaction and re-ranking. GOAnnotator [14] extends this idea by automatically retrieving literature from MEDLINE and retraining a model to re-rank candidate articles using larger datasets, thereby reducing bias and improving robustness. However, such GO-centric approaches are less applicable to viral proteins, because the virus experimental GO annotations are exceedingly sparse [15]. In the UniProt-GOA database (accessed in March 2026), only 0.06% of viral proteins possess experimentally supported GO annotations. This scarcity of labels limits the ability of retrieval-based systems to perform reliable semantic alignment with GO terms. Therefore, direct literature-driven knowledge enrichment becomes a necessary step toward scalable viral protein annotation.

In this context, recent advances in retrieval-based language modeling, particularly retrieval-augmented generation (RAG), have provided powerful methods for integrating external scientific knowledge into reasoning pipelines [16, 17, 18, 19, 20]. A key driver of RAG performance is the quality of the retrieval component: models such as MedCPT [21] jointly optimize semantic retrieval and re-ranking through contrastive pretraining on large-scale PubMed interaction data, effectively bridging user intent with biomedical literature semantics. Despite these advances, applying RAG to viral protein annotation poses unique challenges that existing frameworks have not been designed to address.

The first challenge is viral protein name variability and incomplete literature coverage. Viral proteins are often described under heterogeneous names, aliases, or gene symbols across different studies. Relying exclusively on standardized identifiers, such as those provided by UniProt or RefSeq, often leads to incomplete retrieval of relevant publications. Overcoming this limitation requires retrieval mechanisms capable of recognizing and aggregating diverse mentions of the same protein entity across the literature. The second challenge arises when re-ranking retrieved publications related to viral protein function, as the optimal criteria for prioritizing studies with reliable functional evidence remain unclear. Beyond semantic similarity, additional factors may play important roles, yet their relative importance remains to be systematically examined. A third challenge involves cross-literature knowledge integration. Functional knowledge about a viral protein is typically fragmented across multiple studies that focus on different structural aspects or experimental conditions. Synthesizing such heterogeneous evidence into a coherent functional summary demands multi-stage reasoning that can extract salient facts, maintain contextual consistency, and preserve evidence traceability. Finally, evaluating the quality of enriched function annotations remains an open issue. Traditional classification metrics, such as F1-score or AUROC, are insufficient for measuring annotation enrichment. Evaluation must instead consider semantic correctness, completeness, and evidential grounding, ensuring that generated annotations are not only accurate but also transparent and verifiable.

### Overview

Motivated by these challenges, we propose VirProtRAG, a retrieval-augmented generation (RAG) framework explicitly optimized for viral protein function annotation. As illustrated in Fig. 1, it formulates function characterization as an integrated process comprising three stages: literature retrieval, re-ranking, and evidence-grounded knowledge synthesis. Specifically, for retrieval strategy design, VirProtRAG incorporates LLM-assisted synonym expansion and multi-view, protein-specific query construction, enabling the retrieval system to consolidate diverse aliases describing the same viral protein. Beyond lexical matching, our design leverages MedCPT [21], a PubMed-trained dense retriever, enabling semantic access to contextually relevant literature. In addition, we employ a rank-aware reciprocal rank fusion (RRF) mechanism that adaptively integrates lexical and dense retrieval outputs, providing a more fine-grained fusion than the basic RRF approach. For re-ranking stage, in addition to semantic similarity, we integrate paper-level quality (citation count, venue h-index, author h-index) together with evidence type classification of the target protein, which jointly guide the re-ranking process toward credible and experimentally supported studies. Finally, the top-ranked literature is analyzed using LLMs under a multi-stage prompting scheme that guides the models to understand, filter, extract, and synthesize information, yielding evidence-supported functional descriptions beyond conventional label prediction.

**Fig. 1.**
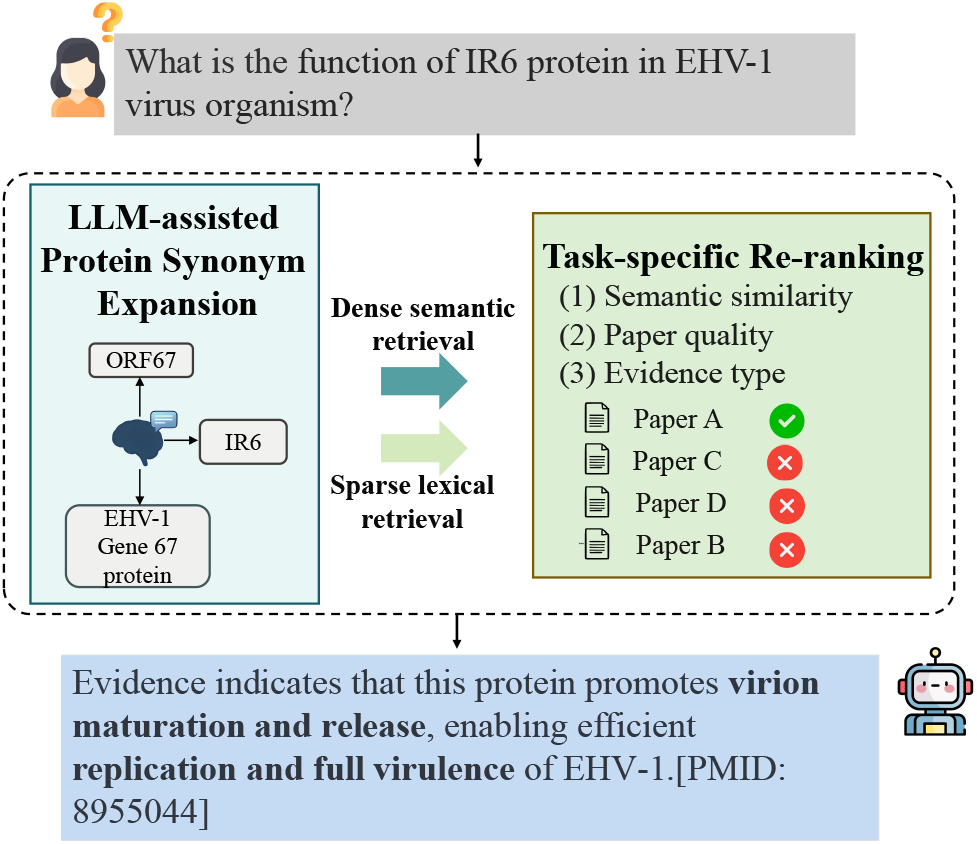
A simplified example of how VirProtRAG applies a task-specific retrieval-augmented generation (RAG) framework to answer user queries by retrieving and synthesizing external knowledge about viral proteins.

Through comprehensive benchmarking against representative retrieval systems, VirProtRAG achieves the highest recall of reference literature. Further analysis attributes this gain primarily to the protein synonym expansion, hybrid retrieval and rank-aware fusion mechanisms. In the subsequent re-ranking phase, the inclusion of paper-level quality indicators and evidence-type awareness leads to consistent improvements across ranking metrics. At the generation stage, seven LLMs from different vendors, covering both open- and closed-source families, were systematically evaluated under expert-provided references (curator-guided), literature retrieved by VirProtRAG (RAG-based), and without any supporting literature (zero-retrieval) scenarios. Overall, GPT-4o consistently achieved the highest semantic alignment with expert annotations and the strongest factual consistency, underscoring its suitability as an intelligent assistant for biocuration workflows. Furthermore, comparisons between the RAG-based and zero-retrieval settings highlight the pivotal role of external knowledge input. Finally, to make VirProtRAG publicly accessible and reusable, we constructed an interactive database containing all 17,484 reviewed Swiss-Prot viral proteins under the Virus taxonomic group (Taxonomy ID: 10239), each annotated by VirProtRAG and searchable by both sequence and text. Among Swiss-Prot annotations originally inferred from sequence similarity rather than direct protein-specific literature evidence, 56.34% could be independently supported by literature retrieved by our system. Case studies demonstrate its ability to enrich uncharacterized viral proteins with literature-linked, biologically meaningful functional insights.

## Methods

### Overview

Given a viral protein with metadata *x* = (*p, g, o*), where *p, g* and *o* denote the protein name, gene name, and organism, respectively, VirProtRAG formulates viral protein function annotation as an evidence-grounded retrieval–generation problem over a biomedical literature corpus *D* (PubMed). Because viral protein nomenclature is lexically diverse, we use an LLM to expand the protein name *p* into a synonym set Syn(*p*) and incorporate this expansion into a multi-view query constructor *Q*(*·*) that maps *x* to a set of queries targeting complementary functional aspects. We then employ a retrieval module *R* to perform hybrid retrieval over *D* and aggregate results into a ranked candidate list, from which the top-*k* papers are selected as the evidence set *E*_*k*_(*x*) = *R*_*k*_(*Q*(*x*); *D*) *⊂ D*. To improve the reliability of downstream generation, the retrieved papers are further re-ranked by reliability-oriented components that prioritize credible and informative evidence, and an LLM generator *G* finally produces a literature-grounded function annotation *ŷ* = *G*(*x, E*_*k*_(*x*)) with explicit links (e.g., PMIDs) to the supporting papers for traceability. The overall architecture is shown in Fig. 2.

**Fig. 2.**
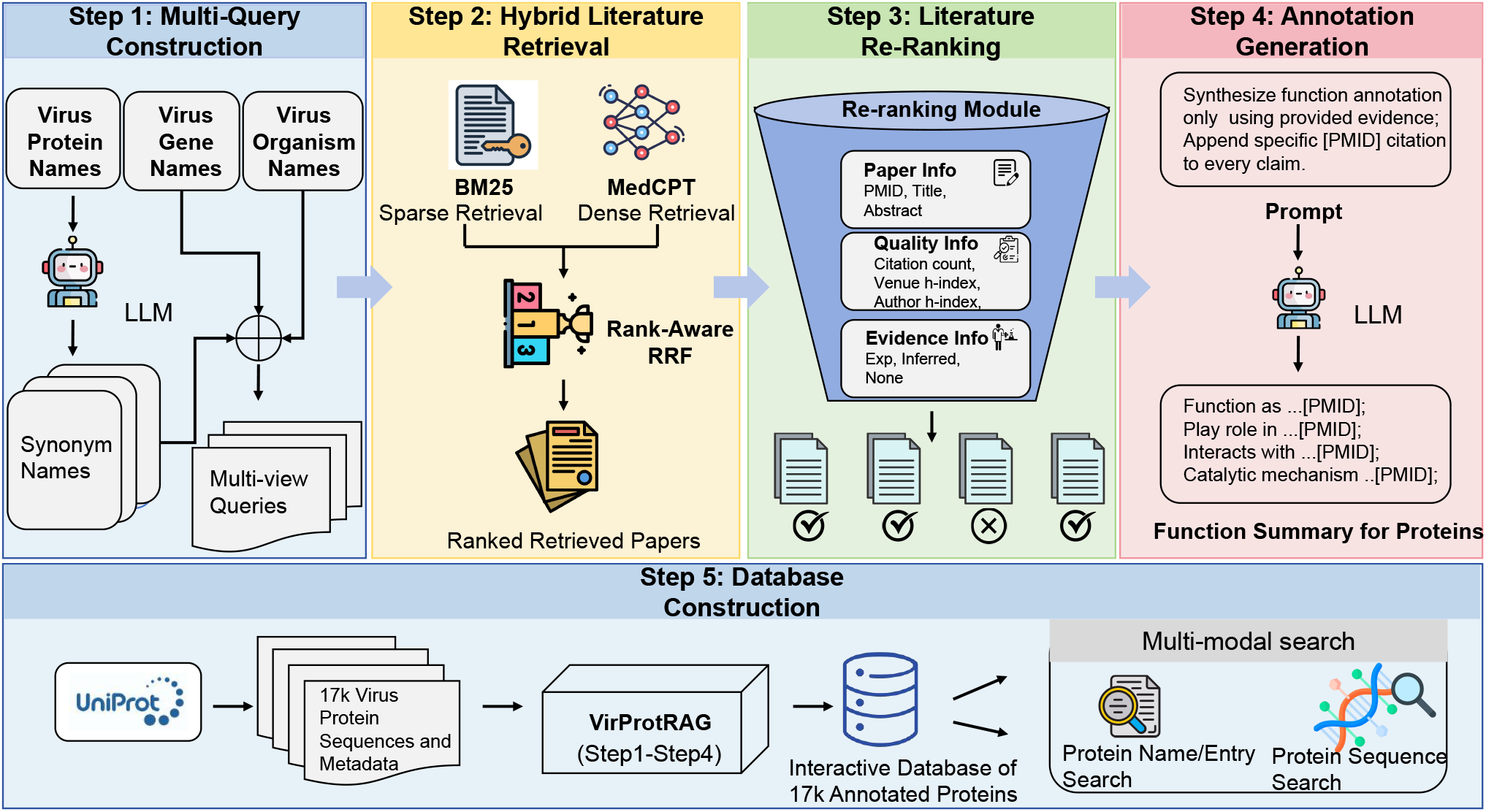
Overview of the VirProtRAG workflow. The pipeline comprises five modules: (Step 1) LLM-based query expansion from protein metadata and multi-view query construction covering complementary functional aspects; (Step 2) hybrid literature retrieval that combines BM25 and MedCPT, followed by rank-aware fusion; (Step 3) reliability-oriented re-ranking that integrates literature signals with quality- and evidence-related features; (Step 4) LLM generation conditioned on the retrieved papers and prompts to produce a literature-grounded functional summary with explicit supporting citations (e.g., PMIDs); and (Step 5) large-scale annotation of viral proteins and construction of an interactive database supporting both text-based and protein-sequence retrieval.

### Two-channel Retrieval

Effective viral protein annotation requires comprehensive and reliable retrieval, covering all potentially relevant literature and avoiding noise accumulation. Guided by this principle, the retrieval module is designed as a two-channel system that integrates complementary search paradigms. It comprises three main components: the corpus, query construction, and retrieval strategies. The corpus is derived from the PubMed database, which contains approximately 39 million bibliographic records and abstracts across the biomedical domain and serves as the primary source for downstream retrieval [22].

As shown in Fig. 2 (Step 1), the queries are constructed based on the protein names, gene names, and organism names. In biomedical text, protein names exhibit substantial lexical variability—the same protein may be denoted by abbreviations or species-specific synonyms. Such heterogeneity often causes conventional keyword retrieval to miss relevant publications [23]. To mitigate this, we employ LLM-assisted synonym expansion using the state-of-the-art model GPT-5 [24].

Beyond lexical heterogeneity, biological functions are inherently multi-dimensional, encompassing molecular mechanisms, catalytic activity, biological processes, and other facets. A single generic query (e.g., “function of protein X”) cannot capture this breadth. Therefore, we design multi-view, biologically motivated query templates integrating the protein name, gene name, and organism name, targeting distinct function aspects. This design enables retrieval to better reflect the multifaceted nature of protein biology.

Building on these constructed queries, we explored both sparse and dense retrieval paradigms shown in Fig. 2 (Step 2). Sparse models leverage exact lexical matching, providing high interpretability and precision, whereas dense models embed queries and documents in a shared semantic space, enabling concept-level matching beyond surface terms. These two channels are thus inherently complementary, one capturing explicit terminology, the other uncovering semantically aligned evidence. Both retrieval channels return the top 1,000 ranked publications by default.

Despite leveraging state-of-the-art LLMs for contextual synonym expansion, occasional inaccuracies or context-inappropriate protein names may arise. Therefore, the retrieval workflow is designed with hierarchical weighting to mitigate such noise. For sparse retrieval, searches initially use the original protein names; synonym-based queries are triggered only when the top-*N* limit has not been reached. For dense retrieval, results from both query types are integrated using a weighted score fusion, where synonym-expanded matches receive slightly lower weights.

To ensure computational efficiency without compromising coverage, we further introduce an adaptive recall-saturation criterion, which automatically determines the optimal cutoff *K*^*∗*^. Given a discrete sequence of recall values {*R*(*K*_*i*_)}, we compute the marginal gain Δ*R/*Δ*K* and identify where additional retrieval becomes uninformative. When the average gain over a sliding window of size *w* falls below a predefined threshold *τ*, the corresponding *K*_*i*_ is set as the saturation point:

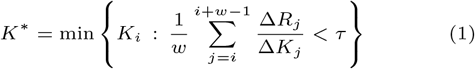

This adaptive cutoff prevents over-retrieval while ensuring that recall remains close to its saturation plateau, thereby yielding an optimal trade-off between computational efficiency and completeness of evidence coverage.

### Rank-aware Reciprocal Rank Fusion (RRF)

After determining the optimal cutoffs *K*_sparse_ and *K*_dense_, the BM25 [25] and MedCPT results are fused using a hybrid approach based on RRF [26], which fully leverages the complementary strengths of sparse and dense retrieval. The core principle of RRF is to assign higher scores to documents that rank highly in multiple lists, relying on their rank positions rather than raw similarity scores, which makes it inherently robust to scale differences across retrieval systems. The basic RRF formula is below:

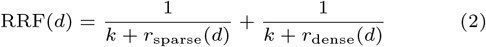

where RRF(*d*) denotes the final fused score for document *d, r*_sparse_(*d*) and *r*_dense_(*d*) denote the rank positions in the sparse and dense retrieval lists, respectively, and *k* is a damping constant (typically 60) controlling the contribution of lower-ranked documents.

This basic RRF employs static and equal weights across different retrieval sources, implicitly assuming that all systems contribute equally at all rank levels. However, empirically, we observe that BM25 tends to outperform MedCPT in higher-ranking positions, while MedCPT gradually surpasses BM25 as the ranking depth increases. Therefore, the conventional RRF introduces two key limitations. First, at shallow ranks, uniform weighting diminishes BM25’s strength in precise lexical matching. Second, at deeper ranks, the same weighting scheme fails to exploit the dense model’s superior semantic recall capability.

To overcome these limitations, we propose a rank-aware RRF that assigns adaptive weights as a function of rank. Specifically, higher weights are allocated to the sparse retriever (BM25) at shallow ranks, where its precision-oriented retrieval is more trustworthy; conversely, the dense retriever (MedCPT) receives increasing weight at deeper ranks, where its semantic generalization improves recall. This rank-dependent weighting mechanism enables the fusion process to dynamically balance precision and recall, producing a more adaptive and theoretically grounded fusion score distribution.

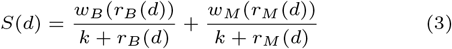

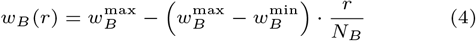

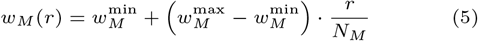

Given two retrieval models, BM25 (B) and MedCPT (M), the final fused score *S*(*d*) for a document *d* is computed using a rank-aware reciprocal rank fusion, as shown in Equations 3–5. Here, *r*_*B*_(*d*) and *r*_*M*_ (*d*) denote the ranks of document *d* returned by BM25 and MedCPT respectively. The functions *w*_*B*_(*r*) and *w*_*M*_ (*r*) define linear rank-dependent weights, where 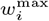 and 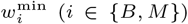 specify the maximum and minimum weighting values, and *N*_*B*_ and *N*_*M*_ represent the number of retrieved candidates from each model. A higher *w*_*i*_(*r*) indicates stronger emphasis on higher-ranked documents, allowing the fusion to dynamically balance the influence of the two retrieval sources.

After hybrid fusion, we again apply post-fusion optimization, the same saturation-based method to determine the best cutoff 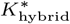 for the combined ranking. The final retrieval results correspond to the top 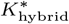 documents, which collectively achieve a high recall while maintaining a manageable candidate size.

### Re-ranking Enhanced by Paper Quality and Evidence Integration

While the hybrid retrieval stage ensures comprehensive recall, additional refinement is required to ensure that the most credible and scientifically valuable papers appear at the top of the final ranking. To this end, the re-ranking stage integrates not only semantic similarity but also two complementary reliability dimensions: paper quality and experimental evidence, which are shown in Fig. 2 (Step 3).

Paper quality reflects a study’s scholarly impact and methodological rigor, serving as an important proxy for information reliability. By introducing a paper-quality feature, the re-ranking model can prioritize well-validated, high-impact research while down-weighting peripheral or less rigorous studies. This reduces the influence of noisy or low-impact publications on the final ranked list.

Complementary to publication quality, the reliability of a function annotation in biological research depends heavily on its evidence source. Experimentally validated findings carry substantially higher confidence than those inferred computationally or lacking explicit support. A system that only considers semantic similarity may inadvertently rank speculative claims above experimentally confirmed results. Incorporating the evidence dimension thus enables preferential ranking of experimentally supported knowledge, improving both scientific interpretability and trustworthiness.

Paper quality is quantitatively modeled as a weighted combination of citation-, venue-, and author-level indicators:

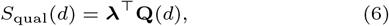

where

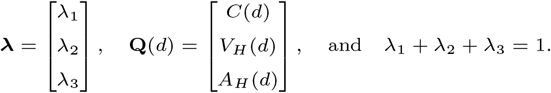

Here, *C*(*d*) denotes the normalized citation count of paper *d*; *V*_*H*_ (*d*) represents the normalized *h*-index of the publication venue; and *A*_*H*_(*d*) denotes the normalized *h*-index of the sum of the first and corresponding author. The weight vector ***λ*** controls the relative contribution of each factor, with the citation component (*λ*_1_) typically dominating due to its strong reflection of research impact. Eq. 6 therefore provides a compact vectorized expression of the overall paper-quality score *S*_qual_(*d*).

To obtain publication-quality indicators, we developed an automated data-collection pipeline that enriches each PubMed record with bibliometric metadata retrieved from the OpenAlex database [27]. The pipeline extracts citation counts, journal-level metrics (e.g., h-index), and author h-indices using local caching, rate-limited API calls, and multi-threaded execution to improve efficiency and reliability.

The evidence feature *S*_evid_(*d*) is derived by prompting LLMs to classify the type of functional support described in each paper abstract into three categories: (1) Experimental – reports direct functional assays or in vitro/in vivo experiments demonstrating the protein’s biological activity; (2) Inferred – suggests potential functions or draws analogies to known proteins without experimental validation; and (3) None – merely mentions the protein without providing functional information. We assign weights of 1.0, 0.5, and 0.0 to these categories, respectively. The detailed prompting design is provided in Supplementary Section 1.

The overall re-ranking score for each document is computed as:

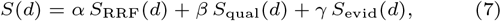

with *α* + *β* + *γ* = 1 and non-negative weights controlling the relative influence of semantic relevance, publication quality, and evidence credibility. Finally, the aggregated score *S*(*d*) is used to rerank all retrieved candidates, ensuring that documents combining strong semantic relevance, high scientific quality, and reliable experimental validation are prioritized in the final output.

## Experimental setup

### Dataset Construction for the Viral Protein Function Augmentation Benchmark

We downloaded all viral protein entries (taxonomy ID: 10239) from UniProtKB/Swiss-Prot. For each entry, we collected the following metadata fields: Entry ID, Protein Names, Gene Names, Gene names (primary), Gene names (synonym), Organism, and the corresponding Function [CC] annotation. Each Function [CC] block consists of one or more free-text sentences, typically accompanied by bibliographic references and evidence codes that specify the provenance and reliability of each functional statement. These codes span a spectrum from direct experimental support (ECO:0000269), author-derived statements in literature without experimental validation (ECO:0000303), and curator inference based on scientific reasoning (ECO:0000305), to sequence-based functional propagation from a characterized homolog (ECO:0000250) or from a curated computational sequence model such as a hidden Markov model profile (ECO:0000255, a formal subclass of ECO:0000250). A detailed description of each code, together with illustrative examples, is provided in Supplementary Table S1.

Each UniProtKB/Swiss-Prot entry is assigned an annotation score on a scale of 1 to 5, reflecting the richness and completeness of its annotation. Stratifying the full viral proteins by annotation score and examining the distribution of evidence codes reveals that the majority of viral protein function annotations (71.46%-80.41%) are derived from sequence similarity (ECO:0000250 and ECO:0000255) rather than from direct literature evidence. The detailed information can be found in Supplementary Fig. 1. Motivated by this observation, and by the distinct nature of the underlying evidence, we partition the data into two complementary datasets as our benchmark datasets, LitBench and SeqBench (Table 1). LitBench contains literature-linked curator annotations and is therefore used for direct performance evaluation of retrieval, re-ranking, and LLM generation. In contrast, SeqBench lacks direct literature references and is used to assess whether VirProtRAG can discover independent literature evidence for annotations originally inferred only from sequence similarity.

**Table 1.** Statistics of annotation scores in the **LitBench** and **SeqBench** datasets.

| Annotation score | LitBench |  | SeqBench |  |
| --- | --- | --- | --- | --- |
|  | #Protein | #Total paper | #Protein | #Function point |
| Score 1 | 43 | 44 | 375 | 383 |
| Score 2 | 507 | 663 | 3621 | 3929 |
| Score 3 | 673 | 1154 | 3464 | 4406 |
| Score 4 | 424 | 916 | 1762 | 2795 |
| Score 5 | 667 | 2719 | 1487 | 12587 |

**LitBench** comprises entries whose Function [CC] annotations are supported by at least one literature-traceable evidence code, including ECO:0000269, ECO:0000303, or ECO:0000305. Each is linked to a PubMed identifier. This dataset serves two purposes: the associated PubMed references are used for benchmarking retrieval and re-ranking performance, and the curated functional text provides an expert-anchored reference for evaluating the quality of LLM-generated annotations.

**SeqBench** comprises entries whose Function [CC] annotations are derived exclusively from sequence similarity (ECO:0000250 and ECO:0000255) and therefore lack direct literature references. For this dataset, our objective is to assess the extent to which VirProtRAG can retrieve primary literature that independently supports these sequence-inferred functional claims. Overall, LitBench contains 2,314 proteins linked to 5,496 PubMed references, whereas SeqBench contains 10,709 proteins with 24,100 sequence-inferred function points.

We note that **LitBench** represents an incomplete subset of all functionally relevant literature, and annotations reflect the current state of established knowledge rather than an exhaustive functional characterization. Consequently, all reported retrieval and generation metrics should be interpreted as lower bounds.

### Evaluation Metrics

To comprehensively assess our retrieval-augmented summarization framework, we evaluate both (i) retrieval and re-ranking effectiveness and (ii) downstream summarization quality. Table 2 summarizes all metrics used in this study, together with their evaluation targets and intuitive interpretations.

**Table 2.** Evaluation metrics for retrieval/re-ranking and downstream functional summarization.

| Metric | Meaning | Brief description |
| --- | --- | --- |
| **Retrieval / Re-ranking metrics** |  |  |
| Recall@K | Retrieval coverage | Fraction of all curator-linked reference literature appearing in the top-$K$ retrieved results (how many relevant items are found). |
| nDCG@K | Ranking quality | How well the top-$K$ ranking promotes relevant literature toward the top, with larger gains for higher-ranked hits (position-discounted, normalized by the ideal ranking). |
| MAP@K | Ranking quality | Measures ranking quality by computing precision each time a relevant item appears in the top-$K$ list, then averaging these values — higher when relevant literature is consistently ranked near the top. |
| **Downstream summarization metrics** |  |  |
| Cosine similarity | Semantic alignment | Embedding-based semantic similarity between the LLM-generated functional summary and the curator-written summary (semantic closeness beyond exact wording). |
| Key Point Recall (KPR) | Content coverage | Fraction of curator-identified key functional points that are present or entailed in the LLM-generated summary (coverage of critical function statements). |
| Faithfulness | Grounding factuality | Fraction of atomic statements in the LLM-generated summary that can be verified or supported by the retrieved evidence (degree of factual grounding). |

#### Retrieval and Re-ranking metrics

We report three standard metrics for retrieval and re-ranking: Recall@K, Normalized Discounted Cumulative Gain (nDCG@K), and Mean Average Precision (MAP@K). All three metrics are computed at the protein level and then macro-averaged across all proteins in the benchmark.

Recall@K measures the proportion of curator-annotated relevant items successfully retrieved within the top-*K*. The nDCG@K is a ranking-sensitive evaluation metric that quantifies how closely a system’s ordering of retrieved documents approximates the ideal ranking of relevant items within the top-k results. Formally, nDCG@K is defined as the ratio between the discounted cumulative gain (DCG) of the obtained ranking and the ideal discounted cumulative gain (IDCG) corresponding to a perfect ranking. In addition, MAP@K considers the precision observed at each rank where relevant literature appears across the top-K retrieval results. A model that retrieves relevant items earlier and with fewer irrelevance yields a higher MAP@K score. The detailed definition for these three metrics is shown in Supplementary Section 2. In addition, the specific example for the calculation is illustrated in Supplementary Fig. 2.

#### Downstream Summarization Metrics

##### Cosine Similarity

To assess the semantic consistency between protein function summaries generated by the LLMs and those curated by human experts, we compute their embedding-based cosine similarity. Both summaries are encoded using the MedCPT query encoder to obtain dense semantic representations.

##### Key Point Recall (KPR)

KPR [28] measures how comprehensively the LLM-generated summaries capture the key functional concepts identified by human curators. To compute this metric, the curator-authored functional descriptions are first decomposed into a set of minimal atomic statements, referred to as key points. The LLM-generated summaries are then segmented in the same way. This segmentation process is implemented through prompting an LLM, which automatically identifies distinct functional statements. Subsequently, another language model compares the two sets and determines, for each curator key point, whether a semantically equivalent or entailed statement appears in the LLM-generated key points. Formally, let *H* = {*h*_1_, *h*_2_, …, *h*_*m*_} denote the set of curator key points, and *M ⊆ H* the subset that are successfully covered by the model-generated key points, where | *·* | denotes the number of a set. The Key Point Recall is defined as:

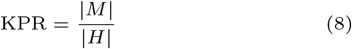

##### Faithfulness

Faithfulness [29] quantifies the factual consistency between an LLM-generated response and the corresponding retrieved or provided contextual sources, which has been employed in the evaluation of RAG systems for various domains, including biomedical text mining and plasmid characterization [30]. It assesses whether each statement in the generated summary can be verified or logically inferred from the evidence contained in the retrieved context. Formally, let the generated response be segmented into a set of minimal factual statements *C* = {*c*_1_, *c*_2_, …, *c*_*n*_}, and let *V ⊆ C* denote the subset of statements verified by the retrieved context. The faithfulness score is defined as:

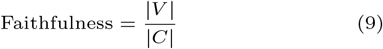

where |*V* | represents the number of verified statements and |*C*| the total number of statements identified in the response. In practice, the evaluation framework RAGAs [29] is used for calculation.

## Experimental results

In this section, we present a comprehensive evaluation of VirProtRAG across multiple experimental dimensions to assess its retrieval, re-ranking, and generation capabilities. First, we compare different retrieval models and strategies to evaluate their effectiveness. Second, we examine the re-ranking enhancements introduced by paper-quality and experimental-evidence features, quantifying their contributions. Third, we evaluate the generation performance of various LLMs under curated, retrieval-augmented, and zero-retrieval settings. Finally, leveraging VirProtRAG, we construct a literature-linked viral protein annotation database, enabling large-scale exploration and downstream analysis. In addition, for existing protein annotations that were originally derived only from sequence similarity, VirProtRAG retrieves relevant publications to provide additional support.

### Comparative Evaluation of Retrieval Models and Strategies

To evaluate the retrieval performance, we first compared four representative methods, including one sparse retriever (BM25) and two dense models (MedCPT and ColBERTv2 [31]), and our proposed hybrid approach VirProtRAG. Fig. 3 shows the Recall@K curves for these methods. Overall, the hybrid model VirProtRAG consistently outperformed both the sparse and dense baselines at all cutoffs. Among dense models, MedCPT performed best, likely benefiting from its biomedical training on PubMed search logs. Notably, BM25 demonstrated superior early precision in the top-ranked results due to its exact term matching when query keywords appeared in titles or abstracts. However, as K increased, its Recall@K plateaued rapidly because lexical matching alone cannot accommodate the diverse terminologies used to describe protein functions. In contrast, MedCPT maintained steady recall growth at larger K, reflecting its ability to retrieve semantically related yet lexically divergent literature. This cross-model comparison establishes VirProtRAG as a strong retrieval backbone for subsequent re-ranking and generation experiments.

**Fig. 3.**
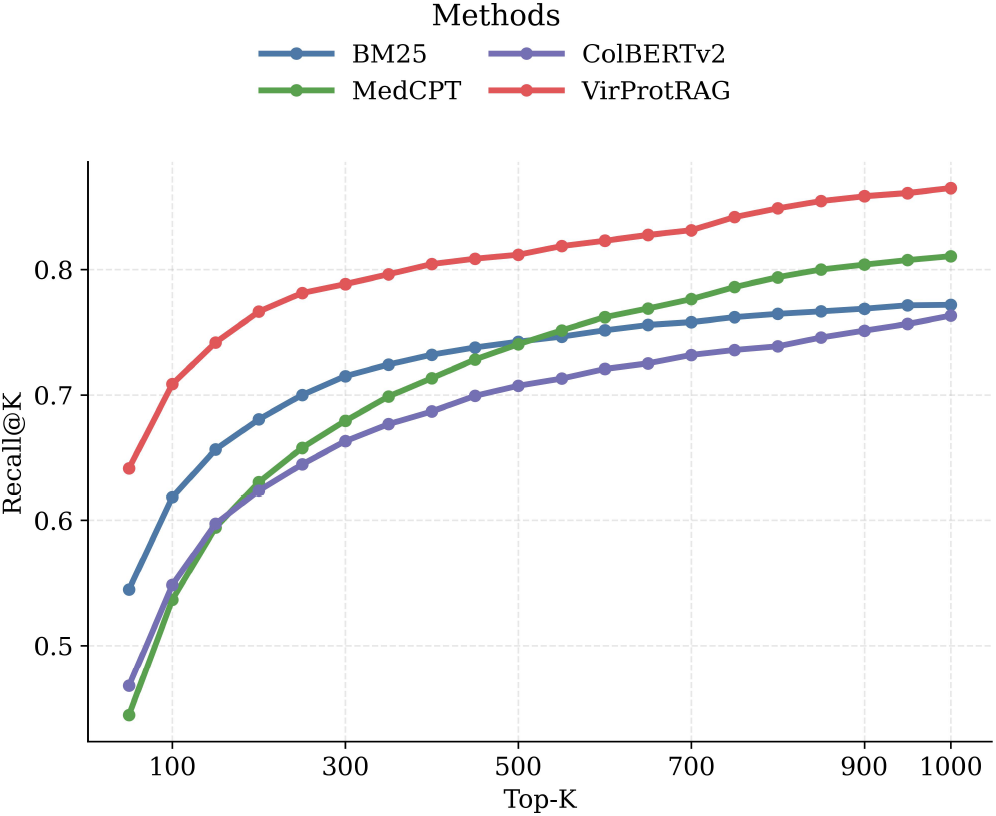
Recall@K comparison across four retrieval methods.

In addition, when evaluating the retrieval recall across different UniProt annotation scores, we observed that literature for proteins with intermediate scores (scores 2, 3, and 4) is generally easier to retrieve. Conversely, the recall for proteins with score 1 and score 5 is significantly lower (Fig. 4). The observed pattern across annotation scores arises from two distinct mechanisms. proteins of Score 1 often have uninformative names—for example, “Protein L”, “32 kDa protein”, “Uncharacterized protein C-102b” —which hinder the retrieval system from constructing effective queries. proteins of Score 5 exhibit low Recall due to two compounding factors. First, Score 5 proteins possess an average of 4.08 PMIDs per entry, substantially higher than Scores 2–4 (1.31–2.16). Identical retrieval quality yields systematically lower Recall for multi-publication proteins. When evaluated using Hit Rate@K (the fraction of proteins with at least one curator-annotated PMID in the top-K results), Score 5 ranks comparably to Scores 2–4 (Supplementary Fig. 3). This confirms that the Recall metric structurally penalizes their richer evidence base. Second, polyproteins account for 34.5% of Score 5 entries, compared to only 0.2–1.2% in Scores 2–4. Within Score 5, polyproteins consistently underperform non-polyprotein entries across all K values (Supplementary Fig. 4). These polyproteins—for example, the SARS-CoV-2 replicase polyprotein that yields 16 mature proteins with 29 supporting references—are annotated with numerous constituent protein names in UniProt. As the number of protein name terms in the query increases, Recall systematically declines (Supplementary Fig. 5). The mechanism is straightforward: broader queries retrieve papers covering all constituent cleavage products, but papers about heavily studied domains (e.g., capsid and envelope proteins) vastly outnumber those about niche functional domains, pushing the latter to lower ranks.

**Fig. 4.**
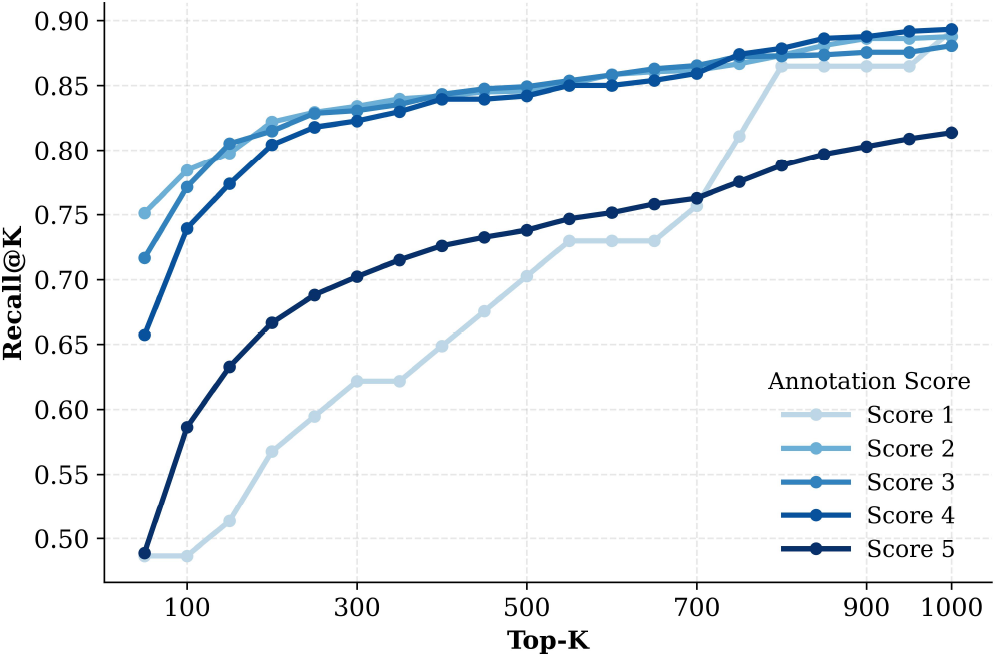
The Recall of top K literature for different annotation scores protein.

To systematically evaluate the contributions of synonym expansion and rank-aware fusion, we evaluate seven retrieval configurations derived from the same underlying BM25 and MedCPT backbone, varying two design choices: the query construction strategy (original protein names only: -baseQ; vs. add LLM-expanded synonyms: -synQ) and, for hybrid methods, the fusion strategy (standard RRF vs. rank-aware variant: - RankRRF). In all cases, the retrieval method itself remains unchanged; only the inputs or fusion differ. The results are shown in Fig. 5.

**Fig. 5.**
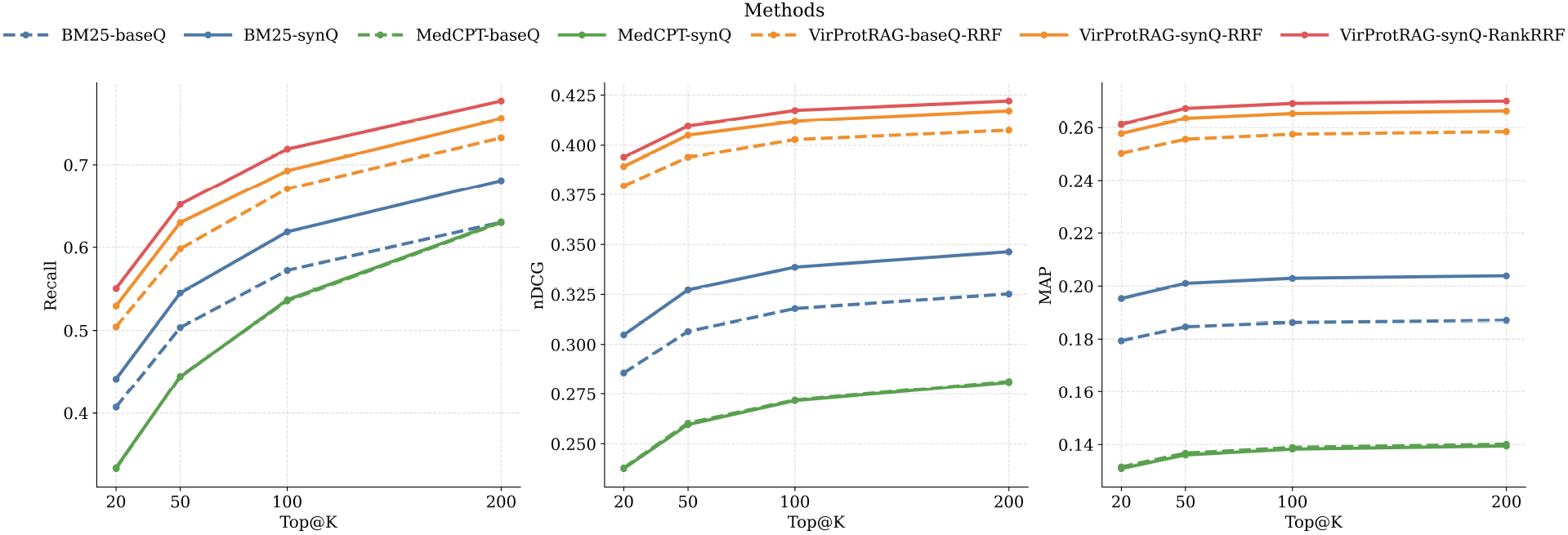
Comparison of seven retrieval configurations evaluated on the viral protein function benchmark. The curves report Recall@K, nDCG@K, and MAP@K across increasing K thresholds. Solid and dashed lines indicate configurations using original (-baseQ) and synonym-expanded (-synQ) queries, respectively. “-RankRRF” denotes our proposed rank-aware fusion variant.

First, comparing BM25-baseQ with BM25-synQ, synonym expansion noticeably improves all three metrics, with the largest gain of 5.05% in Recall@200. This improvement arises because viral proteins are frequently referenced under heterogeneous aliases across publications. In contrast, synonym expansion provides little benefit for MedCPT (MedCPT-baseQ vs. MedCPT-synQ), as dense embedding models inherently encode semantic similarity within their representation space and are therefore less sensitive to surface-level lexical variation. For the hybrid framework, VirProtRAG-baseQ-RRF already achieves competitive performance by fusing BM25 and MedCPT via standard RRF. Introducing synonym-expanded queries (VirProtRAG-synQ-RRF) yields further gains, primarily driven by the enhanced lexical recall of the BM25 component. Finally, adopting our rank-aware RRF (VirProtRAG-synQ-RankRRF) brings additional improvements across all three metrics, with recall increasing by up to 2.62%, through dynamically weighting BM25 at shallow ranks, where its precision is highest, and MedCPT at deeper ranks, where semantic generalization proves more effective.

### Re-ranking Enhancement via Paper Quality and Evidence Features

To assess the contribution of the re-ranking modules, we first identified the optimal retrieval cutoff K=600 based on the saturation point of the recall curve. We then re-ranked the top 600 literature and conducted an ablation study comparing three configurations. VirProtRAG is the complete system, which incorporates paper-quality and evidence-type features into the re-ranking score, and represents our final proposed method. VirProtRAG w/o Evidence additionally excludes evidence-type signals and applies paper-quality-based re-ranking. VirProtRAG w/o re-ranking uses the hybrid retrieval output directly without any re-ranking, serving as the retrieval-only baseline.

As shown in Fig. 6, both additional modules deliver consistent and measurable gains across all metrics in different retrieval depths (K=20, 50, 100, 200). Overall, removing these two feature sets leads to an average decrease of 4.67% in nDCG@K, 4.20% in MAP@K, and 2.52% in Recall@K across all cutoff values. Specifically, excluding experimental-evidence features consistently reduced performance by 1.59%–1.89% in nDCG@K and 2.01%–2.65% in MAP@K. In addition, further removal of paper-quality features results in a decrease in nDCG@K by 2.69%–3.10%, MAP@K by 1.56%–2.18%, and Recall@K by 1.70%–2.45%. The results suggest that both paper-quality and evidence features effectively refine the ranking order. At the same time, the re-ranking stage boosts previously lower-ranked but relevant papers, recovering additional true positives and thereby improving recall.

**Fig. 6.**
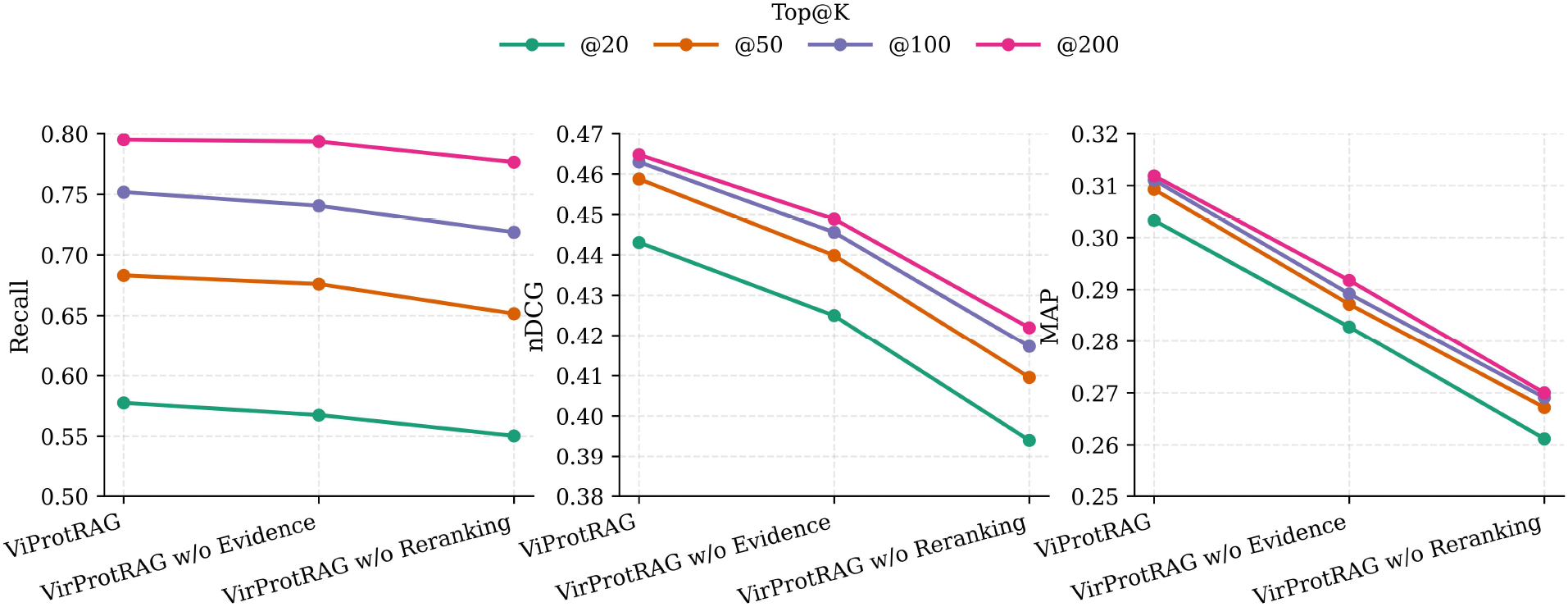
Ablation study on re-ranking components across retrieval depths (K = 20, 50, 100, 200). Three configurations are compared: VirProtRAG (full system), VirProtRAG w/o Evidence (quality re-ranking only without evidence type), and VirProtRAG w/o re-ranking (retrieval only without paper-quality and evidence type).

### Retrieval-based Literature Injection Improves LLM Performance in Protein Function Annotation

Although existing biomedical LLM benchmarks evaluate tasks such as question answering, clinical reasoning, semantic indexing, and literature summarization, they do not directly measure task-specific evidence-grounded functional annotation generation for viral proteins. We therefore evaluated seven LLMs under three settings. Our objective is to understand how both model choice and the source of input affect generation quality in the VirProtRAG pipeline.

#### Scenario 1: Curator-guided annotation generation

In this setting, function annotations are generated based on reference documents human curators have already identified and selected. The LLMs are then guided to extract, synthesize, and summarize the functional information from these curated texts.

#### Scenario 2 :Retrieval-augmented generation (RAG) setting

In this setting, the LLM model receives the top-k documents from the literature retrieval stage as input. Here, we set the top-k to 5, 10, and 20. This configuration mirrors a real-world end-to-end RAG application, where the system automatically retrieves potentially relevant papers and relies on the LLMs to interpret, filter, and generate function annotations directly from this noisy and partially relevant evidence. This scenario tests their robustness to imperfect retrieval results and their ability to discern relevant biological information within a more realistic, less curated environment.

#### Scenario 3 :Zero-Retrieval Settings

In this scenario, the model directly generates protein function annotations without any retrieval input. The LLMs receive only the basic metadata describing the target protein—its name, gene identifier, and source organism. This setup serves as a baseline to evaluate the model’s inherent knowledge and reasoning capability in the absence of external evidence, which helps us to explore the added value of retrieval.

Seven large language models (LLMs) include both closed-source and open-source systems. The closed-source models include GPT-4o-mini [32], Gemini-2.0-Flash [33], GPT-4o [32], and Claude-Sonnet-3.5 [34], while the open-source models include DeepSeek-V3 [35], Qwen2.5-72B-Instruct [36], and GLM-4.7 [37]. To ensure reproducibility and eliminate stochastic variation in the generated results, we set the temperature to 0 to reduce stochastic variation. In addition, to mitigate the computational cost of employing multiple LLMs, we randomly selected 200 proteins for evaluation.

Table 3 presents the results for Scenario 1. Among all evaluated models, GPT-4o achieved the highest KPR (0.7800), outperforming the second-best model (Claude-Sonnet-3.5). For Faithfulness, Claude-Sonnet-3.5 obtained the top score (0.8096), followed closely by GPT-4o (0.8042). With respect to semantic similarity, GLM-4.7 achieved the highest cosine similarity (0.8283), while GPT-4o again ranked second (0.8253). For Scenario 2 (Table 4), increasing the retrieval depth from Top-5 to Top-10 and Top-20 generally improved scores in cosine similarity, KPR, and faithfulness across all models. However, the performance gains from Top-10 to Top-20 were notably smaller than those from Top-5 to Top-10, suggesting a diminishing return effect whereby additional retrieved documents contribute progressively less relevant information. The results of zero-retrieval settings for Scenario 3 (Table 5) show that the semantic similarity and KPR decline for each model compared with the curator-guided annotation generation and RAG setting.

**Table 3.** Generation Performance of LLMs using curator-selected literature (Scenario 1) as input. The best results for each metric are **bolded**, and the second-best results are <u>underlined</u>.

| LLMs | Sim | KPR | Faithfulness |
| --- | --- | --- | --- |
| **Open-source LLMs** |  |  |  |
| Qwen2.5-72B-Instruct | 0.8149 | 0.7351 | 0.7802 |
| DeepSeek-V3 | 0.8206 | 0.7016 | 0.7406 |
| GLM-4.7 | **0.8283** | 0.7426 | 0.7925 |
| **Closed-source LLMs** |  |  |  |
| Gemini-2.0-Flash | 0.8078 | 0.6705 | 0.7875 |
| GPT-4o-mini | 0.8051 | 0.7094 | 0.7613 |
| Claude-Sonnet-3.5 | 0.8244 | <u>0.7516</u> | **0.8096** |
| GPT-4o | <u>0.8253</u> | **0.7800** | <u>0.8042</u> |

**Table 4.** Performance comparison across different retrieval depths (**Top 5, Top 10, Top 20**) between open- and closed-source models using automatically retrieved literature (Scenario 2, retrieval-augmented generation setting). The best results for each metric are **bolded**, and the second-best results are <u>underlined</u>.

| LLMs | Top 5 |  |  | Top 10 |  |  | Top 20 |  |  |
| --- | --- | --- | --- | --- | --- | --- | --- | --- | --- |
|  | Sim | KPR | Faithfulness | Sim | KPR | Faithfulness | Sim | KPR | Faithfulness |
| Open-source LLMs |  |  |  |  |  |  |  |  |  |
| Qwen2.5-72B-Instruct | 0.7678 | 0.6181 | 0.7422 | 0.7833 | 0.6668 | 0.8126 | 0.7925 | 0.7309 | 0.8129 |
| DeepSeek-V3 | 0.7705 | 0.6065 | 0.7511 | 0.7888 | 0.6698 | 0.8144 | 0.7896 | <u>0.7320</u> | 0.8268 |
| GLM-4.7 | 0.7729 | <u>0.6262</u> | 0.7970 | **0.7959** | <u>0.6798</u> | 0.8402 | **0.8057** | 0.7197 | 0.8689 |
| Closed-source LLMs |  |  |  |  |  |  |  |  |  |
| Gemini-2.0-Flash | 0.7479 | 0.5379 | 0.7600 | 0.7754 | 0.6378 | 0.8025 | 0.7810 | 0.6643 | 0.7833 |
| GPT-4o-mini | 0.7615 | 0.6114 | 0.7658 | 0.7825 | 0.6716 | 0.7790 | 0.7844 | 0.7207 | 0.7883 |
| Claude-Sonnet-3.5 | <u>0.7738</u> | 0.6250 | **0.8325** | 0.7829 | 0.6795 | <u>0.8850</u> | <u>0.7964</u> | 0.6953 | **0.8875** |
| GPT-4o | **0.7743** | **0.6416** | <u>0.8212</u> | <u>0.7955</u> | **0.7160** | **0.8945** | 0.7917 | **0.7337** | <u>0.8767</u> |

**Table 5.** Generation Performance of LLMs under Scenario 3 (zero-retrieval setting using only basic protein metadata as input). The best results for each metric are **bolded**, and the second-best results are <u>underlined</u>.

| LLMs | Sim | KPR |
| --- | --- | --- |
| Open-source LLMs |  |  |
| Qwen2.5-72B-Instruct | 0.7455 | 0.5924 |
| DeepSeek-V3 | 0.7369 | 0.5340 |
| GLM-4.7 | **0.7675** | 0.5932 |
| Closed-source LLMs |  |  |
| Gemini-2.0-Flash | 0.7151 | 0.4297 |
| GPT-4o-mini | 0.7443 | 0.5328 |
| Claude-Sonnet-3.5 | <u>0.7570</u> | <u>0.6012</u> |
| GPT-4o | 0.7470 | **0.6109** |

Taken together, these results show that the source and quality of supporting literature have a substantial impact on generation performance. The curator-guided setting generally yielded the best performance. The RAG setting was slightly lower, likely because automatically retrieved documents may include noisy or partially irrelevant evidence that can distract the model during summarization. In addition, retrieved documents may contain relevant information not present in the expert-curated references, potentially reducing overlap with the reference summaries even when the generated annotations are relevant. Nevertheless, RAG consistently outperformed zero-retrieval generation, demonstrating the value of retrieval-based knowledge injection for viral protein annotation. Model choice also affected performance. GPT-4o consistently achieved the highest or near-highest scores across all three metrics. Claude-Sonnet-3.5 showed particular strength in faithfulness, while GLM-4.7 continued to be the strongest open-source model, performing comparably to several closed-source counterparts.

### Database Construction for Literature-Linked Protein Functions

To make literature-grounded viral protein functional knowledge systematically accessible, we constructed a database built upon the VirProtRAG pipeline. The database is designed with two complementary objectives: (i) to provide literature-based evidence for similarity-derived annotations in UniProt that currently lack direct experimental support, as evaluated on the SeqBench dataset; and (ii) to expand the functional coverage of expert-curated annotations by introducing previously uncurated yet biologically relevant function points.

To assess objective (i), we retrieved the top-10 publications for each protein and prompted an LLM to determine whether the retrieved literature substantiates each function point. We define the **Support Rate** as the proportion of function points whose claims are directly verifiable in the retrieved publications. Overall, 56.34% of sequence-inferred function points in SeqBench were independently supported by literature retrieved by VirProtRAG. When stratified by UniProt annotation score, proteins with intermediate scores (2, 3, and 4) achieve high support rates of 62.03%, 75.31%, and 75.92%, respectively, whereas proteins at the two extremes— score 1 and score 5—exhibit substantially lower rates of 39.69% and 28.76%.

To assess objective (ii), we processed all reviewed viral protein entries from UniProtKB/Swiss-Prot (17,484 entries, downloaded in December 2025) through the VirProtRAG pipeline and systematically compared the resulting LLM-derived summaries against existing expert curation. To quantify the proportion of novel, non-overlapping functional terms introduced by VirProtRAG relative to expert-curated annotations, we define the **Gain Rate** as:

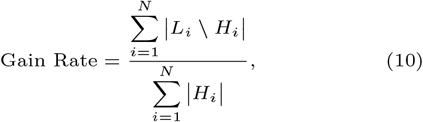

where *L*_*i*_ and *H*_*i*_ denote the sets of LLM-generated and expert-annotated functional points for the *i*-th protein entry, respectively.

Across the entire dataset, VirProtRAG achieved an average Gain Rate of 32.53%, demonstrating that it consistently introduces previously uncurated yet biologically relevant functional knowledge for viral proteins. When stratified by UniProt annotation score, proteins with intermediate scores (2– 4) yield Gain Rates of approximately 43%, whereas proteins with score 1 and score 5 yield substantially lower rates of 13.52% and 17.99%, respectively.

For proteins of score 1, the bottleneck may be literature scarcity: these poorly characterized proteins rarely appear as the primary subject of functional studies, leaving the retrieval pool too sparse to either contribute new function points or substantiate similarity-based annotations. For proteins of score 5, their expert annotations are already saturated, leaving little room for novel function points to be added by RAG and the relatively low support rate may be limited by the challenge in retrieval.

As illustrated in Fig. 7, the database supports multiple retrieval modalities. An amino acid sequence can be submitted for alignment against the internal Swiss-Prot collection using DIAMOND BLASTP [38], returning matched sequences together with their enriched annotations. Alternatively, users can query by protein name, UniProt accession, viral species, or taxonomic identifier to directly obtain the metadata and VirProtRAG-augmented functional descriptions of the corresponding proteins. It provides direct access to the expanded function annotations without the need to re-run the full VirProtRAG workflow, thereby enhancing data accessibility and computational efficiency for downstream analyses.

**Fig. 7.**
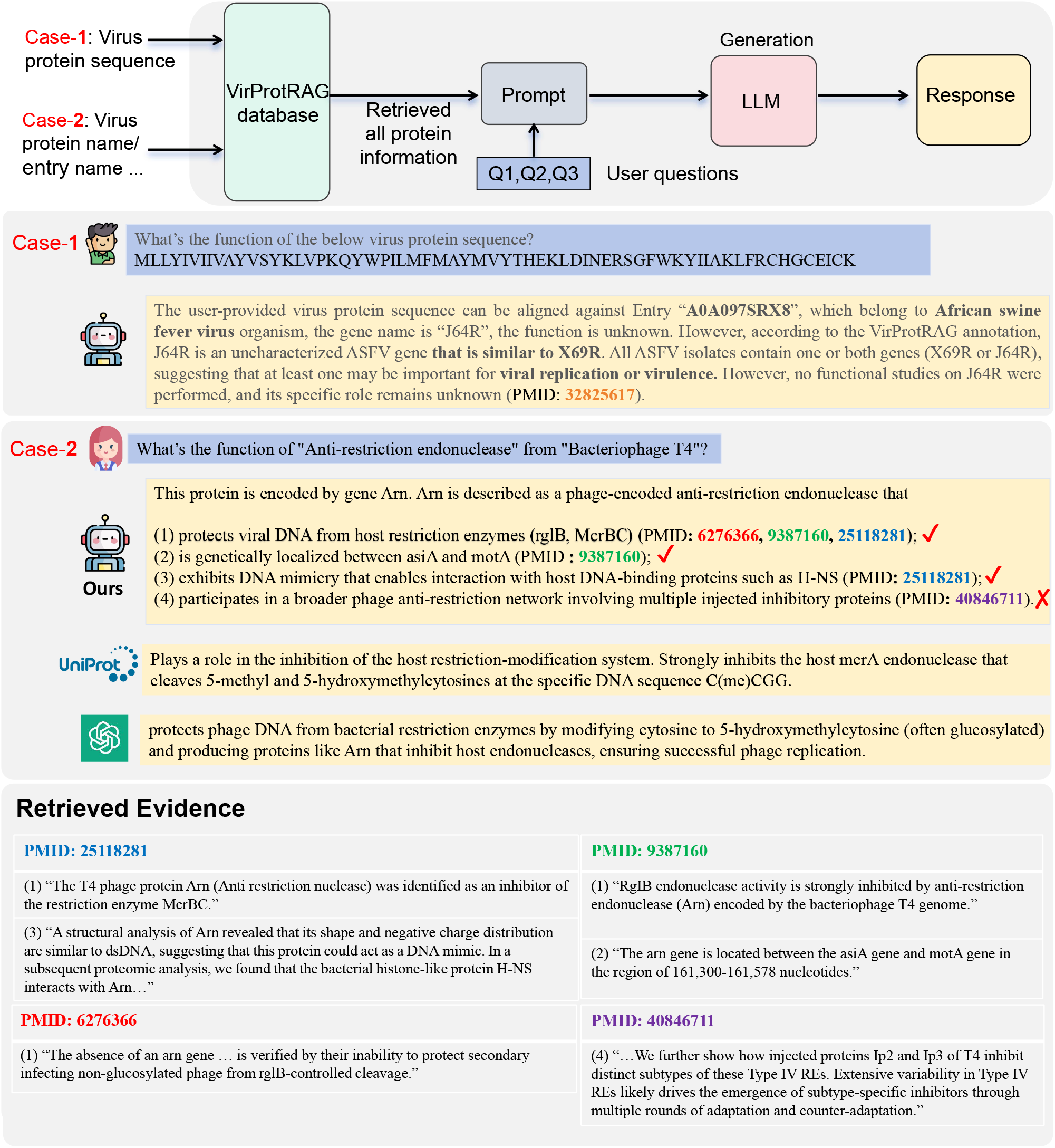
VirProtRAG database query workflow and case studies. The database is constructed using the VirProtRAG-derived protein annotations. Users can query the database either by viral protein sequence (Case-1) or by protein/entry name (Case-2). For each query, an LLM generates a summary. For comparison, we also show the corresponding UniProt annotation and the summary produced by a zero-retrieval (LLM-only) baseline. The retrieved evidence for Case-2 is listed below.

Specifically, when users input the viral protein sequence shown in the Case 1 of Fig. 7, VirProtRAG aligned it to the African swine fever virus (ASFV) gene *J64R*, for which no function annotation is provided in UniProt. It retrieved literature evidence (PMID: 32825617) indicating that *J64R* is an uncharacterized gene closely related to *X69R*. All ASFV isolates contain either *J64R* or *X69R*, implying that at least one of them may play a critical role in viral replication or virulence, although no direct functional studies have been reported. In contrast, fully-LLMs response failed to interpret the protein sequence, producing irrelevant and incorrect answers.

When we query the function of Anti-restriction endonuclease from Bacteriophage T4 shown in Case 2 of Fig. 7, VirProtRAG retrieved four literature-linked statements. Manual inspection of the cited PubMed abstracts confirmed that three were directly supported by cited literature, while the fourth was inconsistent with the referenced study. Compared with the original UniProt description, VirProtRAG captured additional biological details, including the identification of new host restriction targets (*rglB* and *McrBC*) beyond *McrA* mentioned by UniProt, the precise genetic localization of the *Arn* gene, and structural insights revealing DNA-mimicry-based interactions with host nucleoid-associated proteins that facilitate phage infection and resistance to host defense. For the fourth statement, while referring to anti-restriction functions, it described the injected proteins *Ip2* and *Ip3* rather than *Arn*. These proteins inhibit distinct subtypes of Type IV restriction endonucleases. This case illustrates the value of explicit citation links: unsupported or mismatched statements can be rapidly identified through evidence tracing.

Overall, these results highlight that VirProtRAG not only directly accepts the raw protein sequence but also can retrieve relevant supporting literature, producing annotations that go significantly beyond existing database summaries. Moreover, by enriching annotations, the database expands functional coverage for previously uncharacterized and sparse viral proteins and enhances downstream resources such as protein–text alignment corpora and predictive modeling datasets.

## Conclusion and Discussion

In this study, we present VirProtRAG, a novel framework that reformulates viral protein function annotation as an information retrieval and evidence-driven generation problem. The system integrates both lexical keyword–based retrieval and semantic dense retrieval, effectively improving precision and coverage through a rank-aware RRF strategy. By further incorporating paper quality indicators and evidence-type information extracted from the retrieved literature, VirProtRAG performs adaptive re-ranking of literature and substantially advances retrieval reliability and interpretability. Moreover, we developed an interactive and expandable VirProtRAG-derived viral protein annotation database, supporting both protein sequence and text-based queries, which provides an accessible platform for users.

Despite its promising results, VirProtRAG still has several limitations that warrant further investigation. First, the current framework depends on identifiable protein names or gene symbols and associated organisms to construct retrieval queries. For hypothetical or uncharacterized proteins lacking standardized nomenclature in literature or databases, the system cannot effectively retrieve meaningful evidence, reflecting an inherent limitation of text-based retrieval in biology. Second, the system currently relies primarily on abstract-level open-access databases such as PubMed, which limits both the scope and the granularity of accessible knowledge. Certain virus-specific findings are missing, and abstracts often lack detailed experimental descriptions necessary for accurate functional inference. Third, while VirProtRAG integrates sequence-based retrieval through DIAMOND BLASTP, this alignment-based approach remains insufficient for metagenomic datasets containing highly divergent or novel proteins lacking detectable homologs.

In future work, in order to enrich the information space, we aim to incorporate additional open biomedical sources such as bioRxiv, PubMed Central, and other full-text repositories, enabling evidence extraction at the paragraph level and improving both recall and factual grounding. Furthermore, to enhance sequence-based retrieval capabilities for divergent viral proteins, we will investigate deep learning–based protein representation models beyond conventional sequence homology. Finally, although this work focuses on viral proteins, the proposed RAG-based paradigm is generalizable and can be extended to a broad range of microbial domains, such as bacteria and fungi, facilitating large-scale and interpretable function annotation at the microbial community level.

## Supporting information

Supplementary file for VirProtRAG

## Data availability

VirProtRAG is implemented in Python, which can be downloaded from https://github.com/jiaojiaoguan/VirProtRAG.

## Supplementary data

Supplementary Data are available on Zenodo.

## Author Contributions Statement

J.J.G conceived and implemented the algorithms, developed the code, and write the manuscript. J.Y.S. and C.P. reviewed the manuscript and provided constructive suggestions for improvement. Y.N.S. supervised the project, participated in regular discussions, and contributed to manuscript revision. All authors read and approved the final manuscript.

