## Supplementary file for VirProtRAG for "VirProtRAG: Literature-grounded viral protein function annotation with retrieval-augmented generation"

### Supplementary information for "virus protein RAG: comprehensive plasmid characterization and retrieval through sequence-text alignment"

Jiaojiao Guan, Jiayu Shang, Cheng Peng, and Yanni Sun

Electrical Engineering Department, City University of Hong Kong, Kowloon, Hong Kong SAR

April 2026

#### 1 LLM prompt templates for VirProtRAG

The following is the prompt template we used for protein name expansion, function evidence classification, function annotation generation, functional key point extraction, semantic coverage judgment, and literature-based function support identification.

**Prompt Template for Protein Synonym Expansion**

You are a virology and molecular biology expert who is deeply familiar with viral protein nomenclature and synonym usage across databases and academic experimental literature.

**Your task:** Given a JSON input describing an organism and a list of protein names (which may include open reading frame labels, polyprotein cleavage products, or locus tags), identify **reliable synonyms or alternative names** used in scientific papers for each protein *within that organism*.

**Input format (JSON):**

```
{
  "organism_name": "<organism_name>",
  "proteins": ["<protein_name_1>", "<protein_name_2>",
"<protein_name_3>", ...]
}
```

**Output format (JSON):**

```
{
  "synonyms": [
    { "<protein1>": ["<synonym_a>", "<synonym_b>", ...] },
    { "<protein2>": ["<synonym_c>", ...] },
    ...
  ]
}
```

**Synonym inclusion rules:**

1. Include both full and short forms that clearly co-refer to the same protein entity.
2. You may include common descriptive names if they explicitly refer to the same protein in UniProt or PubMed.
3. Do **not** invent speculative names, predicted functions, or inferred homologies.

**Output policy:**

1. Only include synonyms you are **highly confident** are correct for that organism.
2. Do not repeat any protein name that already appears in the input list.
3. If no reliable synonym exists, return an empty list for that protein.
4. Respond using **only valid JSON**, no explanatory text, no Markdown.

##### Prompt Template for Functional Evidence Classification

You are a molecular biology expert specializing in protein function annotation and familiar with UniProt evidence codes.
You will be given the name of a protein and a set of scientific publications (titles and abstracts). For each publication, determine whether it provides *experimental evidence* supporting that protein's biological function, merely *infers* it without experiments, or does not discuss the protein's function at all.
Respond strictly in JSON format as a list of objects, one per paper, like this:

```
[
  {
    "pmid": "123456",
    "label": 0 or 1 or 2,
    "label_name": "NONE" or "INFERRED" or "EXPERIMENTAL",
    "confidence": float between 0 and 1
  },
  ...
]
```

**Label definitions:**

- **0 (NONE):** No protein/gene function described.
- **1 (INFERRED):** Similar ECO code ECO:0000303. Function is *predicted* or *implied* (no experimental validation).
- **2 (EXPERIMENTAL):** Similar ECO code ECO:0000269. The study contains experiments directly testing or demonstrating protein function (e.g., enzymatic assays, mutagenesis affecting activity, knockout/overexpression altering phenotype, etc.).

Respond using **only valid JSON**, no explanatory text, no Markdown.

##### Prompt Template for Functional Annotation Generation

You are a professional **molecular virologist and protein biochemist** responsible for curating UniProt FUNCTION annotations.
Your goal is to read and reason through retrieved publications and produce factually correct, well-evidenced FUNCTION statements for the target protein only.

**TASK CONTEXT**
You will receive:

- Basic protein metadata: **gene names**, **protein names**, and **organism source**.
- A set of **retrieved publications**, each with a **PubMed ID**, **title**, and **abstract**.

These papers may vary in relevance — some are about the target protein, others may be unrelated. Your task is to **filter, analyze, and extract** only biological function information directly about the input protein.

**GENERAL PRINCIPLES**

1. **Scientific Accuracy First**
   - Use only statements explicitly supported by the provided texts.
   - Never infer beyond what is stated in the title or abstract.
   - If unsure whether a claim is supported, exclude it.
2. **Citation Integrity**
   - Every functional claim must be traceable to at least one PubMed ID in the input.
   - Each cited PubMed ID must genuinely contain or directly support the statement.
   - Never fabricate or link a PubMed ID not describing that function.
3. **No Hallucination**
   - Do not invent data, experiments, or functions.
   - Do not assume similarity or homology implies function.
   - Summarize only what the evidence texts actually demonstrate.
4. **Reading Discipline**
   - Process publications one by one, sequentially.
   - After reading each, summarize its findings before moving on.
   - Do not mix evidence across papers until the integration step.

**STEP-BY-STEP WORKFLOW**
**Step 1 — Parse Input**

- Extract all PubMed IDs under `"publications"`.
- Note the target protein's gene names, protein names, and organism.
- Treat each publication independently.

**Step 2 — Assess Relevance (Filtering)**

- **Relevant:** The title/abstract clearly investigates the target protein (exact name or synonym) in the specified organism, and reports biologically meaningful information.
- **Irrelevant:** The paper does not focus on this protein in this organism or only offers generic mentions.
- Keep only relevant papers; discard irrelevant ones.

**Step 3 — Sequential Extraction (Per-Paper Reading)**

- For each relevant publication:
  1. Read title and abstract carefully for factual statements about the target protein's role.
  2. Identify experimental findings describing its function or biological role.
  3. Ignore unrelated methods or entities.
  4. Record (a) concise functional statement, (b) supporting PubMed ID.
  5. Skip papers without function-related content.

**Step 4 — Integrate Findings**

- Merge statements describing the same biological role.
- Keep distinct functions (e.g., "entry", "fusion", "immune evasion") separate.
- Combine relevant PubMed IDs.

**Step 5 — Write Final FUNCTION Lines**

- Format each integrated function as: `FUNCTION: [concise factual statement] (PubMed:XXXXXXX, PubMed:YYYYYYY).`
- Use UniProt tone: concise, objective, mechanistic.
- Recommended verbs: *Functions as*, *Mediates*, *Promotes*, *Inhibits*, *Facilitates*, *Coordinates*, *Required for*, *Plays a role in*, etc.

**Output JSON Format**

```
{
  "Summary": "FUNCTION: ... (PubMed:YYYYYY, PubMed:ZZZZZZ). ...
(PubMed:AAAAAA)."
}
```

If no publication provides any relevant content:

```
{
  "Summary": "No supported functional information found among retrieved
publications."
}
```

**Important:** Do not include markdown, code fences, or extraneous commentary. Return only the final JSON object as output.

#### Prompt Template for Functional Key Point Extraction

You are a biomedical domain expert specializing in protein annotation. Your task is to carefully split a given protein FUNCTION description into a list of concise, independent key points.
**Each key point must:**

1. Represent one independent biological function or action of the protein.
2. Contain a complete and self-contained statement (subject + verb/action + object).
3. Keep conditional or contextual phrases when needed.
4. Preserve uncertainty cues like "may" or "possibly".
5. Avoid merging multiple verbs into a single point.

**Output format (JSON):**

```
{
  "entry": "<entry_id>",
  "key_points": [
    "First key point sentence.",
    "Second key point sentence."
  ]
}
```

**Prompt Template for Semantic Coverage Judgment**

You are a biomedical domain expert in protein function annotation. Your job is to judge the semantic coverage between two sets of key points describing protein functions.

**Task:** For each protein entry, you will be given:

- A list of **Human** key points, labeled as H1, H2, H3, ...
- A list of **LLM** key points, labeled as L1, L2, L3, ...

You must decide which LLM key points semantically *cover* which Human key points.

**Definition of "cover":**

- "Cover" means that the LLM key point conveys the same main biological meaning as the Human key point — even if the wording or phrasing differs.
- Small stylistic or contextual differences (e.g., oxidative stress, localization) can be ignored as long as the core function or mechanism is equivalent.
- If an LLM key point is more general but logically includes the Human key point's meaning, consider it covered.
- If the main function or biological action is missing, consider it *not covered*.

**Output requirements:**

- A single LLM point (e.g., L1) can cover multiple Human points (e.g., H1 and H2), or none.
- Similarly, a Human point can be covered by multiple LLM points if they share equivalent meaning.
- Also list:
  - which Human points are **not covered** by any LLM point, and
  - which LLM points are **extra** (do not match any Human point).

**Output format (JSON):**

```
{
  "entry": "<ENTRY_ID>",
  "coverage_mapping": [
    {"llm": "L1", "human": ["H1"]},
    {"llm": "L2", "human": ["H2", "H3"]}
  ],
  "uncovered_human_points": ["H4"],
  "extra_llm_points": ["L3","L4"]
}
```

**Style constraints:**

- Do not rewrite or explain any sentences.
- Do not output text outside the JSON object.
- Ensure the JSON format is syntactically valid.
- Use the given H# and L# identifiers exactly as presented.

**Prompt Template for Literature-based Function Support Identification**

You are an expert biocurator. Your task is to identify which candidate publications provide evidence supporting specific protein function points, and to characterize that evidence rigorously.

**Inputs:**

1. Target protein metadata: gene name(s), protein name(s), and organism name(s).
2. A numbered list of function points (claims to be verified).
3. A list of candidate publications, each with PMID, title, and abstract.

**Core Principles (READ CAREFULLY):**

- Be **conservative**: it is better to return no support than to fabricate one.
- A publication supports a function point **only if** the title or abstract contains an explicit textual span that directly entails or strongly implies the claim. Vague topical relevance is **not** support.
- You **must** quote the exact supporting span from the abstract or title. If you cannot quote a concrete span, do not list the PMID.
- Verify that the publication truly concerns the target protein (matching gene/protein name) or a clearly stated homolog. Do not assume.

**Evidence Types (ECO codes):**

- **ECO:0000269** — Direct experimental evidence on the target protein in the specified organism (e.g., the abstract reports an experiment demonstrating this function).
- **ECO:0000303** — Non-traceable author statement / review: the paper asserts the function but does not describe original experiments.
- **ECO:0000250** — Sequence similarity / homology: evidence is from a homologous protein in another species; inferred for the target by similarity.
- **ECO:0000305** — Curator inference: the claim is supported only by combining partial information across statements, not by a single explicit assertion.

**Output Format (JSON):**

```
{
  "1": [
    { "pmid": "9557680", "eco": "ECO:0000269", "quote": "exact
verbatim span copied from the abstract or title" }
  ],
  "2": [],
  "3": [
    { "pmid": "10400780", "eco": "ECO:0000250", "quote": "..." }
  ]
}
```

**Hard Rules:**

- If no publication provides explicit support for a function point, return an empty list `[]` for that key. Do not invent support.
- The `"quote"` field must be a verbatim substring of the provided title or abstract — do not paraphrase.
- Do not include any text outside the JSON object (no markdown, no commentary).

#### 2 Retrieval and Re-ranking metrics

**Recall@K** Recall@K is a classical metric that measures the proportion of ground-truth relevant items successfully retrieved within the top-$K$, respectively.

$$\text{Recall@K} = \frac{\sum_{i=1}^{K} rel_i}{|R|} \tag{1}$$

where $R$ denotes the set of all ground-truth retrieval literature for the given query, and $|R|$ is the total number. $i$ denotes the rank position. Each retrieved document $d_i$ is assigned a binary relevance label $rel_i \in \{0,1\}$, where $rel_i = 1$ denotes that the document matches a ground-truth reference for the given query, and $rel_i = 0$ otherwise. A higher $Recall@K$ indicates a more extensive coverage of the relevant collection.

**Normalized Discounted Cumulative Gain (nDCG@K)** The nDCG@K is a ranking-sensitive evaluation metric that quantifies how closely a system's ordering of retrieved documents approximates the ideal ranking of relevant items within the top-k results. Higher nDCG@K values indicate that relevant documents appear closer to the top of the ranking, whereas lower values imply that relevant items are positioned deeper in the list or are missed entirely.

Formally, nDCG@K is defined as the ratio between the discounted cumulative gain (DCG) of the system-produced ranking and the ideal discounted cumulative gain (IDCG) that corresponds to the best possible ordering.

$$\text{nDCG@K} = \frac{\text{DCG@K}}{\text{IDCG@K}} \tag{2}$$

The DCG@K aggregates the relevance of all retrieved documents with a logarithmic position-based discount:

$$\text{DCG@K} = \sum_{i=1}^{K} \frac{2^{rel_i} - 1}{\log_2(i+1)} \tag{3}$$

$(2^{rel_i} - 1)$ ensures that higher relevance levels contribute more strongly, while $\log_2(i+1)$ reduces the impact of lower-ranked documents, emphasizing correct ordering near the top of the list. To obtain the ideal score, we compute the same expression after sorting the documents by descending relevance, where $rel_i^*$ represents the relevance values in the ideal ranking.

$$\text{IDCG@K} = \sum_{i=1}^{K} \frac{2^{rel_i^*} - 1}{\log_2(i+1)} \tag{4}$$

**Mean Average Precision (MAP@K)** The MAP@K measures the consistency of a system's precision across the top- K retrieval results. It integrates both the number of relevant documents retrieved and the precision observed at each rank where a relevant item appears. A system that retrieves relevant items earlier and with fewer irrelevance yields a higher MAP@K score.

The Average Precision at K (AP@K) for a query is defined as:

$$\text{AP@K} = \frac{1}{\min(K, |R|)} \sum_{k=1}^{K} \text{Precision@k} \cdot rel_k \tag{5}$$

where Precision@k is the precision computed at cutoff $k$, and the summation accumulates precision values only at ranks where relevant documents occur. The normalization by $\min(K, |R|)$ prevents an artificial suppression of the score when the number of ground-truth documents exceeds the cutoff depth $K$. To evaluate performance across multiple protein queries, MAP@K averages the per-query AP@K values:

$$\text{MAP@K} = \frac{1}{Q} \sum_{q=1}^{Q} \text{AP@K}_q \tag{6}$$

where $Q$ is the total number of protein queries and $\text{AP@K}_q$ denotes the average precision for query $q$.

### 3 Detailed Evidence code description

UniProtKB/Swiss-Prot employs five primary evidence codes to characterize the reliability of functional annotations in the virus protein function description. ECO:0000269 denotes annotations directly supported by published experimental evidence, with the corresponding PubMed identifier provided for traceability. ECO:0000303 applies to annotations derived from author statements in scientific articles without experimental validation. ECO:0000305 represents annotations inferred by a curator through scientific reasoning, either from domain expertise or from the broader content of a publication.

The remaining two codes capture sequence-based inference. ECO:0000250 (sequence similarity evidence used in manual assertion) is applied when a function is propagated from a specific experimentally characterized protein via pairwise sequence similarity, with the source UniProt accession provided as reference. ECO:0000255 (match to sequence model evidence used in manual assertion) is a subclass (is_a) of ECO:0000250 in the Evidence and Conclusion Ontology, and applies when the annotation is supported by a match to a curated computational model, such as a hidden Markov model (HMM) profile or a HAMAP rule (e.g., ECO:0000255|HAMAP-Rule:MF_04067).

**Supplementary Table 1.** Summary of UniProtKB/Swiss-Prot evidence codes describing the reliability and provenance of functional annotations.

| Evidence Code | Full Name | Evidence Basis | Reference Format |
|---|---|---|---|
| ECO:0000269 | Experimental evidence | Published experimental results directly demonstrating protein function. | PubMed ID |
| ECO:0000303 | Non-traceable author statement | Author statement in a publication without direct experimental support. | PubMed ID |
| ECO:0000305 | Curator inference | Annotation inferred by a curator from domain expertise or from the scientific content of a publication. | PubMed ID (if article-based) |
| ECO:0000250 | Sequence similarity evidence | Function propagated from an experimentally characterized protein via pairwise sequence similarity. | UniProt accession of source protein |
| ECO:0000255 | Match to sequence model evidence | Function inferred from a curated computational sequence model (e.g., HMM profile, HAMAP rule); *subclass of ECO:0000250*. | Model or rule identifier (e.g., HAMAP) |

### 4 Example of Retrieval and Re-ranking Evaluation Metric Calculation

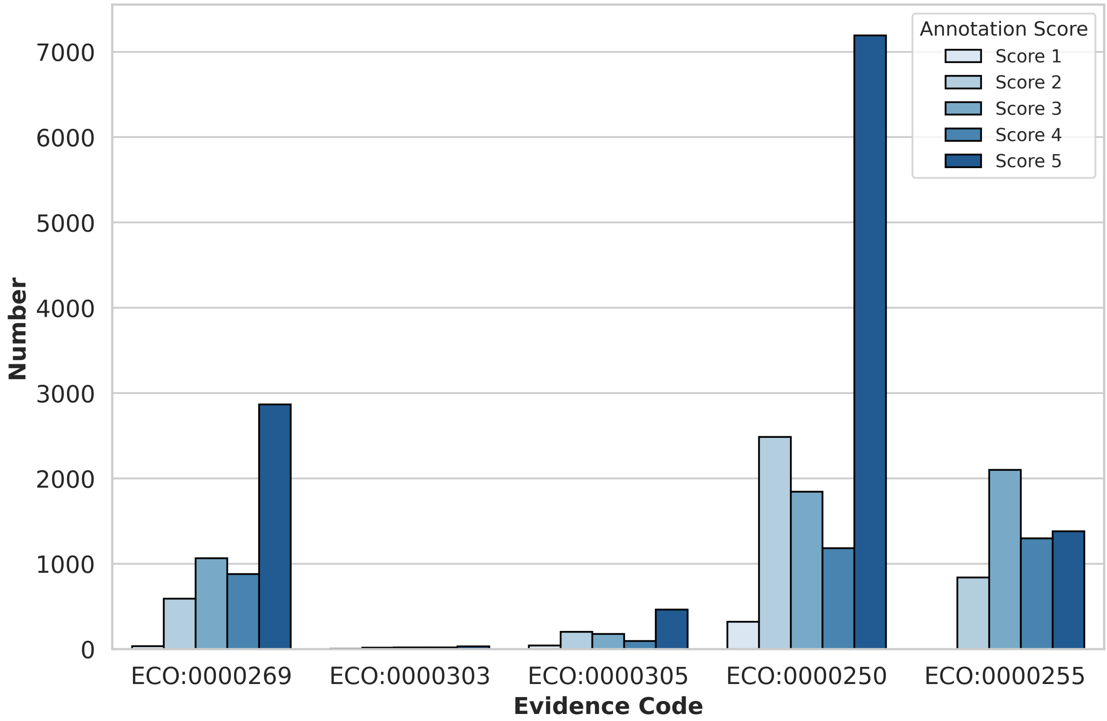


**Supplementary Figure 1.** The distribution of the evidence code for different annotation score.

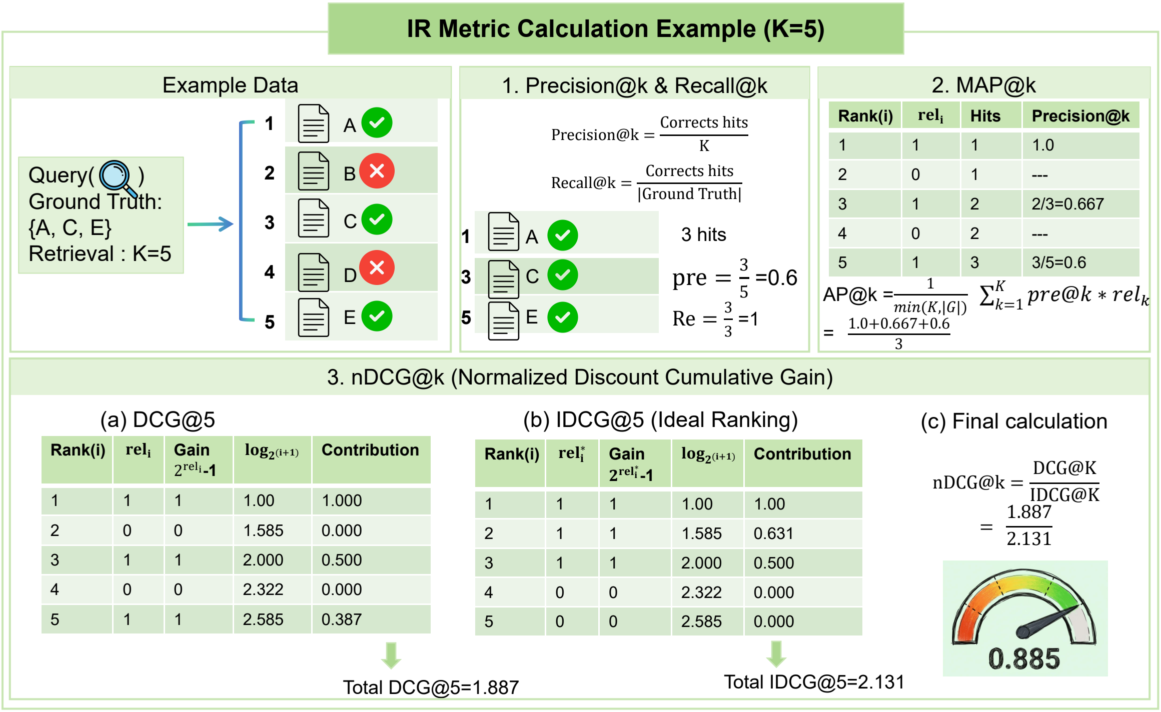


**Supplementary Figure 2.** Illustrative example of retrieval and re-ranking evaluation metric calculation. The example demonstrates the computation of information-retrieval metrics with a retrieval depth of K=5, including Precision@K, Recall@K, MAP@K, and nDCG@K. Example data (left) shows retrieved items A–E and the ground-truth set A, C, E. The upper panels detail how Precision, Recall, and MAP are derived from ranked results, while the lower panels illustrate DCG and IDCG calculations and their normalization to obtain nDCG.

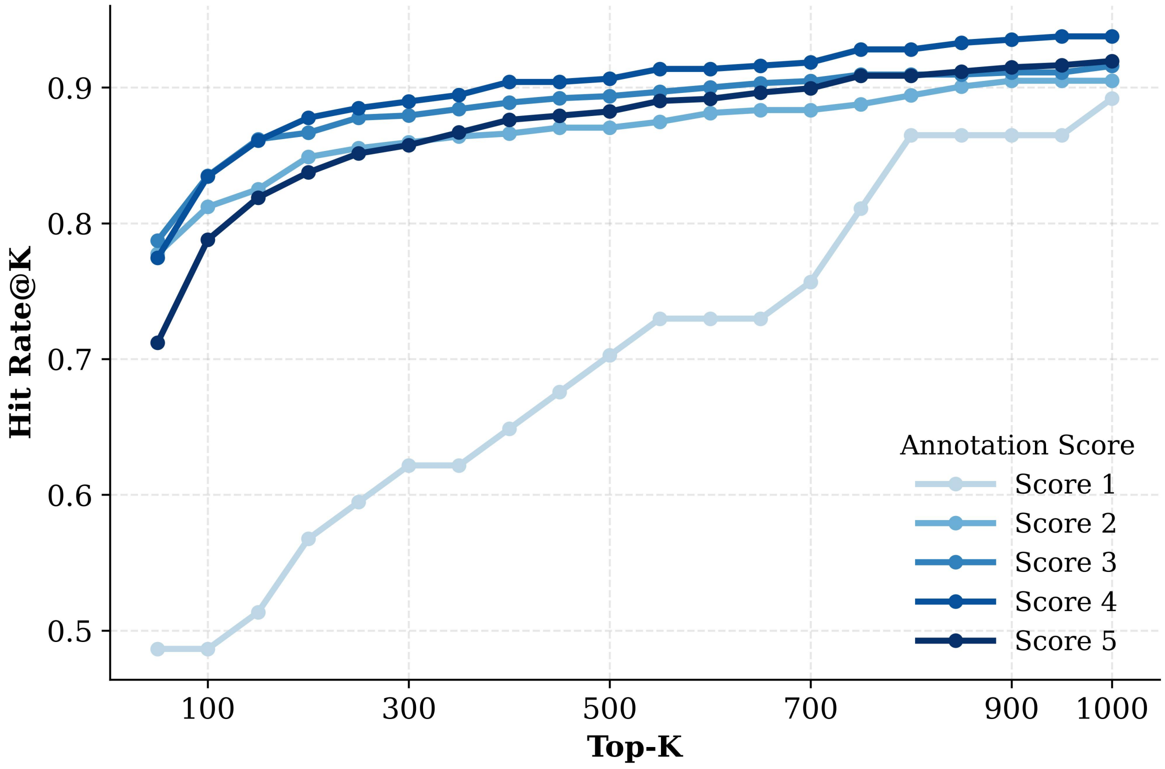


**Supplementary Figure 3.** Hit Rate@K (the fraction of proteins with at least one gold PMID retrieved in the top-K results across annotation scores.

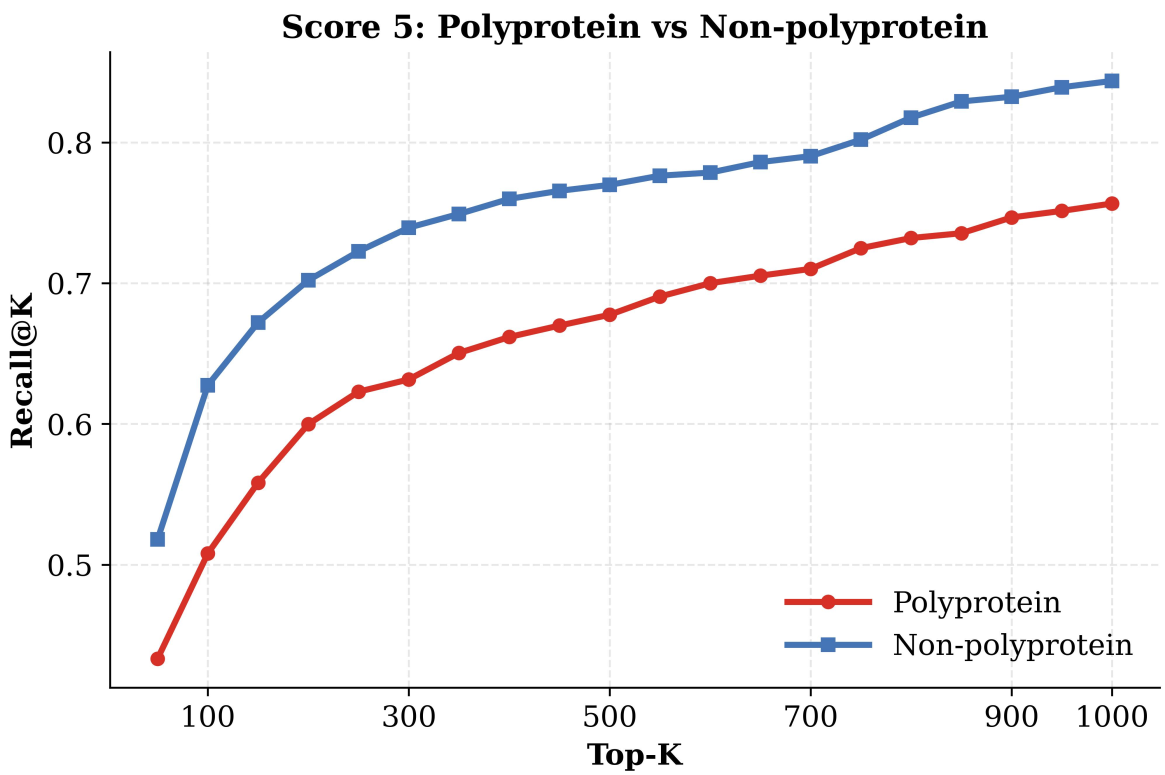


**Supplementary Figure 4.** Recall@K comparison between polyprotein and non-polyprotein entries within annotation Score 5.

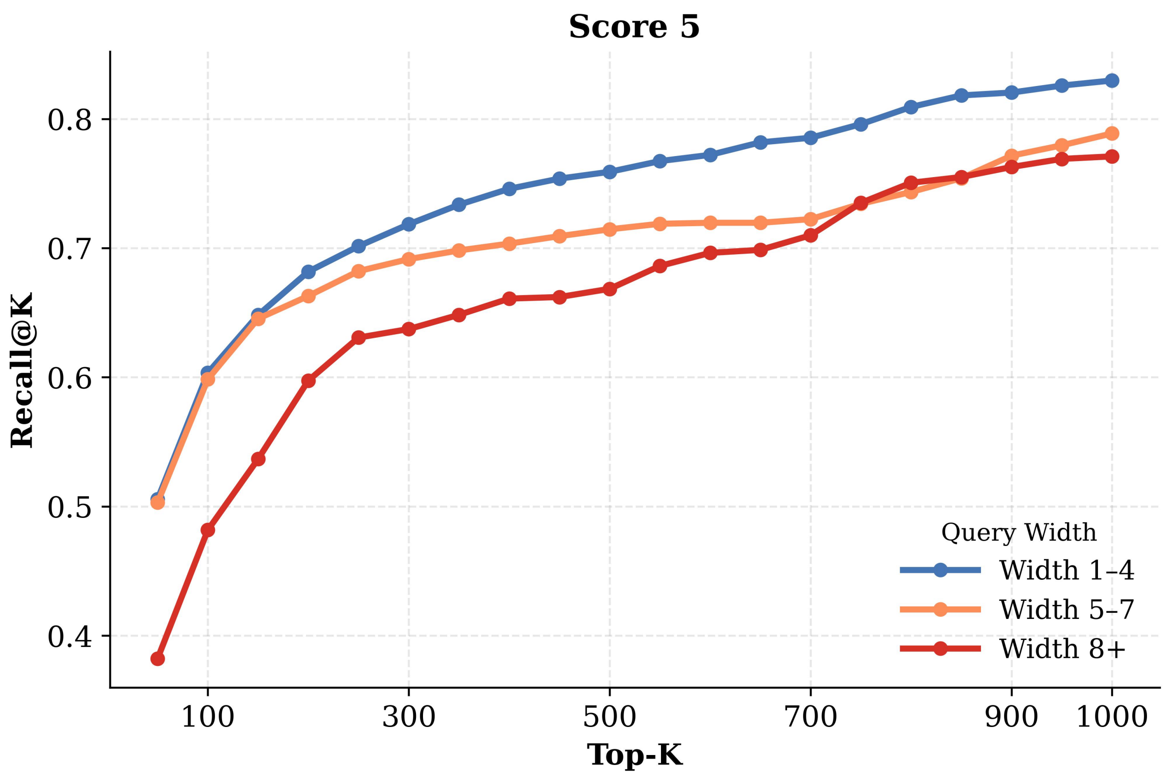


**Supplementary Figure 5.** Recall@K for Score 5 entries stratified by query width (number of protein name terms).
